# Vessel recruitment and derecruitment mechanism and paradoxical behaviour of vasoconstriction during ischaemia

**DOI:** 10.1101/045864

**Authors:** Kayalvizhi Lakshmanan, Sarangam Majumdar

## Abstract

In this paper, we introduce mathematical aspect of recruitment and derecruitment mechanism of blood vessels acting in such a way as to preserve physiological values of the capillary pressure and perspective of pressure drop effect which has been pointed out by a series of studies documenting a paradoxical vasoconstriction during ischaemia.

## I. Vessel recruitment and derecruitment mechanism

Perhaps the only point in which all the current theories and experiment of circulatory system are found to be in agreement is the time-dependent nature of the blood vessels. It therefore seems of considerable interest to investigate the temporal behavior of the number of blood vessels in the microcirculation. It is true that there exits a fairly large number of literature where experimental biologist and doctor study has been made on similar problems; however they all try to explain this phenomenon based on experimental data. The blood circulatory system is a pressure dependent phenomenon which supplies sufficient oxygen and nutrients to every tissue in the body. This oxygen and nutrients rich blood are delivered to the surrounding tissues by the up stream artery(A), large arterioles(LA), small arterioles(SA) and capillary(C). On the other hand, capillaries(C), small venules(SV), large venules(LV), downstream vessels(V) are carrying deoxygenated blood. Heart is the pumping station and connected by up and down stream vessels[1,2]. Depending upon the diameter of the vessel we can divide all the ves Department of Information Engineering, Computer Science and Mathematics, University of L’Aquila, Italy sels into seven different types.Fig.1 illustrates the seven segment model for circulation.

**Fig. 1.**
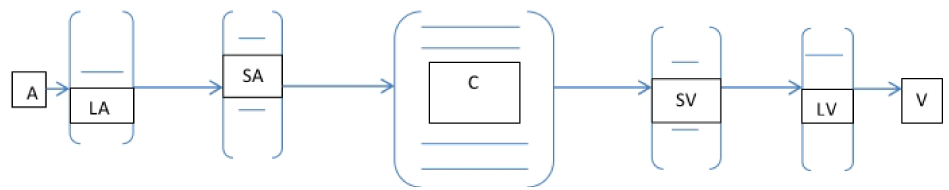
Seven segment model and the direction of the flow.

In the past few years, most of the models try to investigate the vasomotion (spontaneous oscillation inside the vessel). Theoretical model have been used to explain this natural phenomenon to predict the condition of vasomotion using potential mechanism[3,4]. Arciero and Secomb proposed a model [5] on spontaneous oscillations in a model for active control of microvessel diameters. They investigate the dynamics of vasomotion considering diameter and activation both are time-dependent.From Arciero-Secomb model, we can derive first order differential equation for number of vessels which is varying with time in recruitment and derecruitment mechanism as follows.

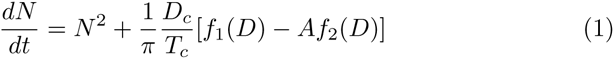

where A is the degree of Vascular smooth muscle activation or tone, *f*_1_(*D*) = *KF*_1_(*D*) and *f*_2_ = *KF*_2_(*D*), 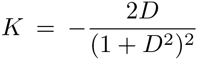 is exponential function as[5].

We assume *F*_2_(*D*) is a Lorentzian function. Here *D* is the diameter of the vessels, *D*_*c*_ and *T*_*c*_ are reference values of the diameter and tension. Fig.2 illustrates that number of vessel(*N*) is increasing with time and this number is quite large enough inside the capillary.

**Fig. 2.**
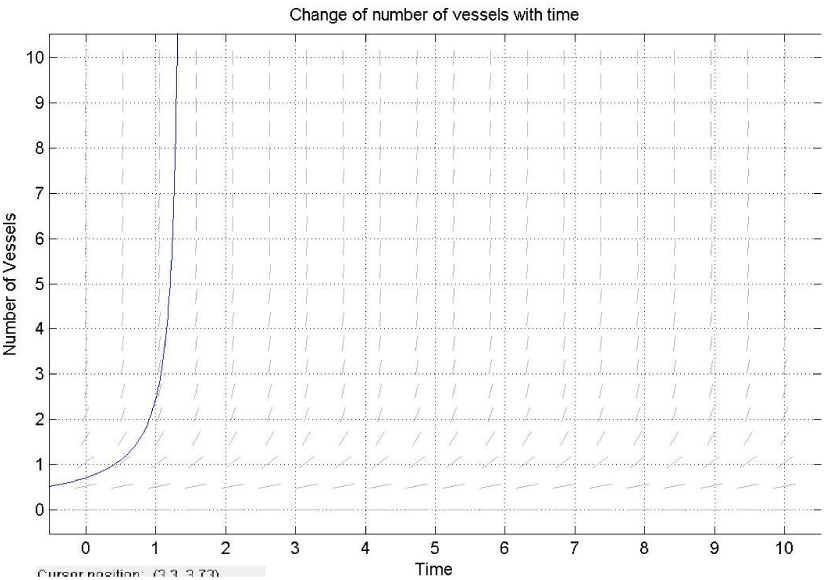
Number of Vessels are changing with respect to time.

From the experimental evidence this number of vessel are related with the diameter. Table 1 shows the changes of diameter and the number of vessels from one segment to another segment and pressure drop and the resistance of all the segments.

**Table 1.** Diameter (*D*) is given in *μ*m, Segment resistance (*R*) in *mmHg.cm*^3^.*s*(×10^8^) and Pressure drop (Δ*P*) is considered in *mmHg*. *N* is number of vessels in each segments and *R* is Segment resistance[6].

| Description | A | LA | SA | C | SV | LV | V |
| --- | --- | --- | --- | --- | --- | --- | --- |
| $D$ | - | 65.2 | 14.8 | 6 | 23.5 | 119.1 | - |
| $N$ | - | 1.00 | 143 | 5515 | 143 | 1.00 | - |
| $R$ | 1.88 | 2.81 | 7.50 | 2.81 | 0.86 | 0.28 | 0.19 |
| $\Delta P$ | 10 | 15 | 40 | 15 | 4.60 | 1.49 | 1.00 |

The spontaneous oscillation inside the blood vessel is pointed out theoretically and experimentally. So we can think about the origin and the effect of noise in the microcirculation which is an open field for research.

## II. Paradoxical behaviour of vasoconstriction during ischaemia

In this section, we are trying to catch the attention of the reader into one of the main heart disease in the western countries. This disease is known as myocardial ischaemia which is one of the cause for heart attack. Ischamia is a vascular disease due to insufficient arterial blood flow to tissue. It can be caused by atherosclerotic plaques in coronary arteries. Most of the models of coronary physiology assume a microvascular adaptation (vasodilation) to the upstream flow resistance that preserves as much as possible coronary blood flow. Vasodilation is a process of widening of blood vessels. But there is one opposite process which is known as vasoconstriction. Vasoconstriction means the narrowing of the blood vessels resulting from contraction of the muscular wall of the vessels, in particular the large arteries and small arterioles. In the recent study there are many clinical articles which are devoted to vasconstriction as a main cause for ischamia.The diameter of the vessels are narrow due to vasconstriction which dirctly implies the blood flow is decrease at that time.Classical formula tells us Δ*P* = *QR* where *Q* is blood flow rate. So the blood pressure in increase at that moment which is one of the cause for stroke. This behaviour is not normal behaviour. In the sense of normal behaviour, we see that blood pressure is decreasing with the number of vessel is increasing in circulation. This kind of behaviour is pointed out in recent studies. They call it as a paradoxical behaviour. However, Doppler monitoring of coronary blood flow (CBF) documented severe microcirculatory vasoconstriction during pacing-induced ischemia in patients with coronary artery disease. In the year 2005, some researchers in Italy researched in this same field and reported that the increase in coronary microvascular resistance observed in patients with coronary artery disease during ischemia is due to the exclusion of a tissue fraction from perfusion. This phenomenon might reflect either vascular collapse due to compression of endocardial layers by endocavitary pressure or by active vasoconstriction. Moreover, this heterogeneously distributed vasoconstriction might actually reflect intrinsic vascular control of vasomotor tone tuned to preserve stable values of microvascular pressure in a selected number of vascular units[7,8].

## III. Conclusions

Demand of tissue varies according to some factors like infection, injury, growth and metabolism. These factors play an important role in our circulatory system. Recruitment and derecruitment mechanism is connected with this paradoxical behaviour in microcirculation which is challenging task for researcher for further investigation.

